# FALCON-meta: a method to infer metagenomic composition of ancient DNA

**DOI:** 10.1101/267179

**Authors:** Diogo Pratas, Armando J. Pinho, Raquel M. Silva, João M. O. S. Rodrigues, Morteza Hosseini, Tânia Caetano, Paulo J. S. G. Ferreira

## Abstract

The general approaches to detect and quantify metagenomic sample composition are based on the alignment of the reads, according to an existing database containing reference microbial sequences. However, without proper parameterization, these methods are not suitable for ancient DNA. Quantifying somewhat dissimilar sequences by alignment methods is problematic, due to the need of fine-tuned thresholds, considering relaxed edit distances and the consequent increase of computational cost. Additionally, the choice of the thresholds poses the problem of how to quantify similarity without producing overestimated measures. We propose FALCON-meta, a compression-based method to infer metagenomic composition of next-generation sequencing samples. This unsupervised alignment-free method runs efficiently on FASTQ samples. FALCON-meta quickly learns how to give importance to the models that cooperate to predict similarity, incorporating parallelism and flexibility for multiple hardware characteristics. It shows substantial identification capabilities in ancient DNA without overestimation. In one of the examples, we found and authenticated an ancient *Pseudomonas* bacteria in a Mammoth mitogenome.

FALCON-meta can be accessed at https://github.com/pratas/falcon.

## INTRODUCTION

In metagenomics, DNA from several organisms are sequenced simultaneously. Using efficient anti-contamination protocols, metagenomics allows looking into ancestral samples for determining composition, such as ancient pathogens. Among other goals, identifying ancient pathogens may help in inferring ancestral death causes ^1^, predicting microbial communities ^2, 3^ and filtering incorrect information in genome assembly ^4^.

Sequencing archaeological remains from the neolithic and paleolithic period is a very difficult process, mostly because of the low quantity and quality of DNA present in the samples, which are associated with the high degree of substitutions, generally caused by PCR amplifications ^5^ and post-mortem degradation ^6^. Also, there are strong computational challenges, such as: dealing with a large volume of raw data with very short read sizes, uncertainty in the degree of contamination of the ancient DNA (aDNA) samples ^7^, very unbalanced composition in terms of sample sizes, lack of a complete catalog of extant and extinct species, lack of knowledge of the degree of dissimilarity between ancestral and present-days species.

The identification of metagenomic composition in ancestral samples has been mostly addressed using small- and large-scale approaches. On a small-scale, the use of 16S rRNA gene sequences has been the most common genetic marker, namely because its sequence is conserved ^2, 8^. However, in aDNA, the current methods have distinguishability problems and ambiguity on similar organisms, mostly because PCR amplification biases can confound standard metabarcoding analyses ^9^.

The general approaches to detect and quantify metagenomic sample composition are based on the alignment of the reads, according to an existing database containing reference microbial sequences ^10^–^16^. Probably, the most used are BWA ^16^, MUMMER ^13^, BOWTIE ^15^ and MEGAN ^14^.

Without proper parameterization these methods are not suitable for aDNA ^17^. Quantifying somewhat dissimilar sequences by alignment methods is usually problematic, due to the need of fine-tuned thresholds, considering relaxed edit distances and the consequent increase of computational cost. Aware of this need, the PALEOMIX ^18^ and MALT ^19^ methods have been proposed using the BWA ^16^ and MEGAN 14 alignment algorithms, respectively, with custom parameters for metagenomic ancient genomes.

However, the choice of the thresholds poses the problem of how to quantify similarity without producing overestimated measures (**Supplementary Note 4**). As such, due to the uncertainty associated with the data from the ancient samples, there is the need for fast probabilistic methods capable of producing conservative similarity estimates. Such methods are those based on the estimation of the amount of information given by compression algorithms ^20, 21^, a fast, revolutionary and efficient way to measure the relative similarity between multiple sequences, including non-assembled aDNA sequences.

In this paper, we present FALCON-meta, a compression-based method to infer metagenomic sample composition of aDNA. It shows substantial identification capabilities without overestimation. It is an alignment-free method that deals with small and large-scale, namely complete human genomes from FASTQ samples.

## METHODS

Appropriate data compressors can be used to estimate similarity between sequences. First, a model is built using reference data. These reference data can be obtained from a sequence or from a collection of fragments (FASTQ reads). Then, the model is kept static, i.e., it is frozen, and the target data is compressed using exclusively the information contained in the reference model. This restriction implies that the compressor will not be able to explore intra-target redundancies. We refer to this type of compression as *relative compression*.

The efficiency of the compressor depends on how well it collects and organizes the information contained in the reference data, so that questions about the target can be quickly and efficiently answered (i.e., with the fewest bits as possible). If a target is efficiently described by the reference, then its relative information is approximately zero. Contrarily, if the target is totally unrelated, it needs the maximum number of bits to describe the target.

We define the Normalized Relative Similarity (NRS) using relative compression and the mathematical concepts provided in **Supplementary Note 1**. Accordingly, the method computes the NRS of each sequence in the database (FASTA) relatively to the whole ancient reads (FASTQ). The top NRS values are stored in a cache (**Supplementary Note 2**).

Choosing efficient models is very important to compute the relative compression. Our implementation uses soft blending of Markov models with several orders. It has been shown that these models, working in cooperation, are very powerful, being superior to state-of-the-art alignment methods in handling very small shuffled reads containing multiple mutations ^21^. Moreover, we combine them with substitutional tolerant Markov models (**Supplementary Note 1**), which are particularly useful to handle the specific characteristics of aDNA, mainly the large amount of substitutions.

The implementation (FALCON-meta) directly works with data from sequencers, independently of the coverage used. This tool can use large sequence databases of known organisms, such as the entire viral, bacterial, archaeal and fungi databases, although it can also run with non-assembled sequences. FALCON-meta uses efficient data structures, namely cache-hashes, and supports multi-threading. The **Supplementary Note 12** describes how to install and use the FALCON-meta tool. The tool can be accessed at https://github.com/pratas/falcon

## RESULTS

Evaluation using synthetic data (**Supplementary Note 3**) shows that alignment-based methods, overestimate similarity (**Fig. 1a,b**) and, therefore, originate a large number of false positives. On the other hand, FALCON-meta does not overestimate similarity. Regarding time complexity, alignment-based methods spent increasingly more time as the number of reads increase, in the synthetic dataset (**Fig. 1c**). On a real scenario (**Fig. 1d,e**), using BOWTIE and BWA with appropriate parameters set for aDNA ^17^, aligning the Neanderthal reads with only one *Escherichia coli* took approximately the same time as FALCON-meta in the entire database (viruses, bacteria, archaea and fungi).

**Figure 1:**
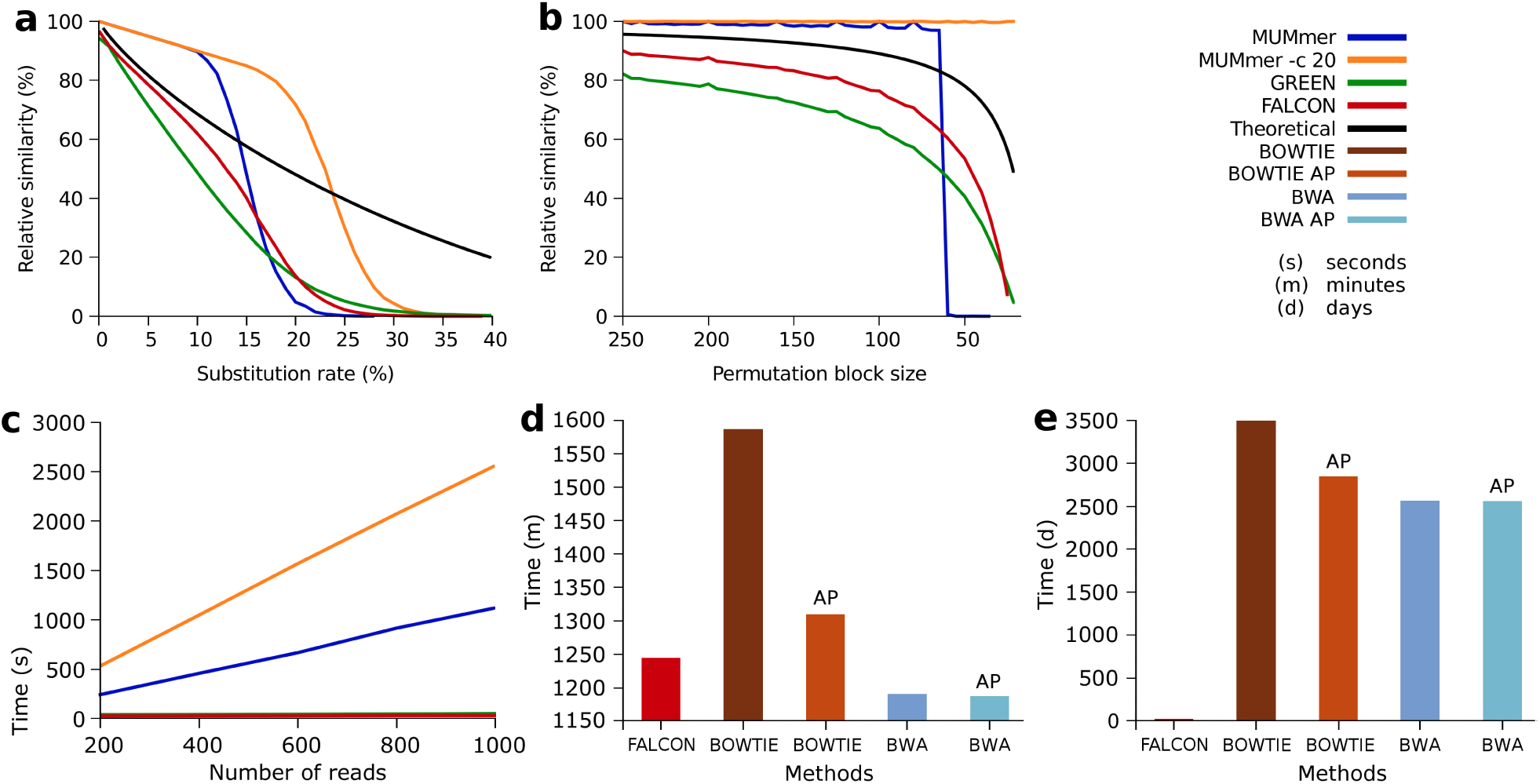
Evaluation of the methods using synthetic and real data. (**a**) Relative similarity of several methods (MUMmer ^13^, MUMmer -c 20 ^13^, GREEN ^20^ and FALCON-meta) compared with the theoretical value, for increasing substitutional random mutation rates. Similarity values above the theoretical line are considered overestimation (**Supplementary Note 4**). (**b**) Relative similarity of the methods (used in (a)), along with the theoretical value, for increasing sizes of random permuted blocks. (**c**) Time spent by several methods (the same as in (a)), for increasing sizes of reads. (**d**) Time spent, in a real case scenario, by the methods (FALCON-meta, BOWTIE ^15^, BOWTIE AP ^15^, BWA 16 and BWA AP ^16^) in the detection of an *Escherichia coli* in the full Neanderthal FASTQ samples. (**e**) Time spent, in a real case scenario, by the methods used in (d) while mapping 3,170 reference organisms in the full Neanderthal FASTQ samples. The time for FALCON-meta is empirical, while for the others methods is an estimation (given the huge amount of time initially spent). See **Supplementary Note 3** for more information. AP stands for customized parameters for ancient DNA.

We have used FALCON-meta on the Neanderthal (**Supplementary Note 7**) and Denisova (**Supplementary Note 8**) high coverage genomes to measure similarity against multiple databases from NCBI, including viruses, bacteria, archaea and fungi. Details on how the experiences have been built, including resources, parameters and the computing environment used to run FALCON-meta are provided in **Supplementary Note 2**.

We have found multiple reference sequences sharing high degree of similarity relatively to the Neanderthal samples, including viruses, bacteria and fungi. Multiple bacteria have been detected, namely *Shigella* spp., *Escherichia coli, Propionibacterium acnes, Clostridium perfringens, Acinetobacter johnsonii*, among others. All the *Shigella* species have been detected (*S. boydii, S. sonnei, S. flexnery* and *S. dysenteriae*).

The identification of metagenomic sample composition must be carefully analyzed, mainly because of the database inter-similarity. For example, the inter-similarity between the *Shigella* spp. and *E. coli* bacteria is very high (**Supplementary Note 5**), therefore, the assumption of both being present in the samples must be carefully considered.

For viral sequences, we have found human endogeneous retrovirus and several phages, such as dsDNA *Enterobacteria phage cdtI* and *Propionibacterium phage Enoki*. Their known hosts are *Escherichia coli* (*Enterobacteriaceae* family) and *Cutibacterium acnes*, respectively. Therefore, these have a high probability of being present on the samples since both genomes and respective phages sequences have high NRS values. More precisely, *Propionibacterium acnes* and *Enterobacteria kobei*.

Several fungi have also high NRS values, through their mtDNA, namely *Penicillium polonicum, Penicillium roqueforti, Pseudogymnoascus pannorum, Zasmidium cellare, Aspergillus flavus*, among others. The first two are from the same genus, thus, it is possible to see a high degree of inter-similarity when *Penicillium polonicum* is measured relatively to *Penicillium roqueforti* (**Supplementary Note 7**). Therefore, the indication of *Penicillium roqueforti* in **Fig. 2** seems to be given by sharing similarity with *Penicillium polonicum*. Moreover, when its local similarity is aligned with the visual map from NCBI (**Supplementary Note 10**), it shows an uniform distribution pattern along with the sequence, unlike in the Denisova genome.

**Figure 2:**
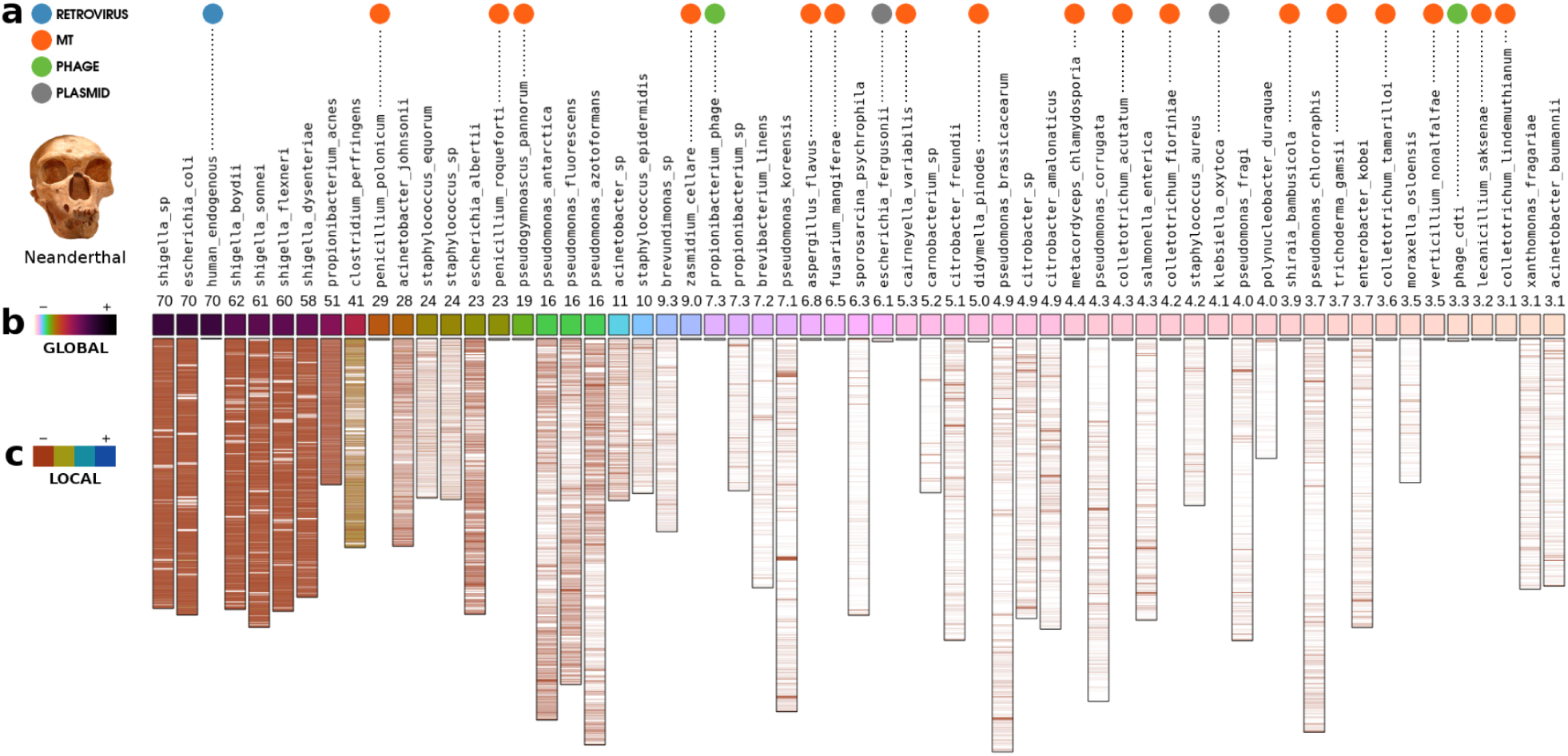
Metagenomic sample composition of a Neanderthal genome raw data sequences inferred by FALCON-meta. (**a**) Classification of the complete genomes, where the absense of circle stands for bacterial genomes. The MT is mitochondrial DNA. (**b**) Relative similarity between a Neanderthal whole genome, using the FASTQ samples, and multiple NCBI databases (for relative similarities *>* 3). (**c**) Local similarity of filtered regions with relative similarity above 0.5 (**Supplementary Note 10**).

The metagenomic sample composition of the Denisova (**Supplementary Fig. 8**) shows also that human endogenus retrovirus sequences are strongly present. In general, the NRS value is lower as the common ancestor is older, for example is lower for chimpanzee and gorilla, and higher for modern humans (**Supplementary Note 6**).

Multiple fungi sequences have been detected in the Denisova samples, such as *Penicillium polonicum, Penicillium roqueforti* and *Pseudogymnoascus pannorum*. Also several bacteria were detected, namely *Escherichia, Shigella, Staphylococcus* and *Clostridium* (**Supplementary Note 8**). When analyzing the results, considering the similarity between the genomes database, we found several organisms that share similarity, namely *Shigella* and *Escherichia* genus. Several species from these genera appear with the highest NRS, because they share high similarity with other organisms from the database.

Increasing the computational resources to search specific viral squences in the Denisova genome we found high similarity with the human adenovirus C with at least 7 % of NRS (**Supplementary Note 10**). When looking for the local similarity of human adenovirus C, relative to the Denisova genome, and aligning with the NCBI visual map, we found that the similarity is located in the L1, L2 and the beginning of the L3 genes.

The identification of a virus similar to the Human Adenovirus C sugests that the Denisova individual was a young child. However, since the virus may persist for some years following the infection ^22^, although unlikely, the Denisova may had been older. These genetic insights are in accordance with the Denisova bone analysis, where the bone growth plates and size led to a prediction of an age between 6 and 13.5 years ^23^.

Additionally, we have inferred the composition of a Mammoth mitogenome ^24^. For example, we have found a sequence of a *Pseudomonas* bateria with similarity to *Pseudomonas antarctica*, a bacteria that has been isolated in Antarctica, with high resistance to low temperature environments (**Supplementary Note 9**). After, we have autheticated the *Pseudomonas* as ancient (**Supplementary Note 11**) and dismissed the possibility of similarity between the mtDNA of the Mammoth and bacterial DNA ^25^.

## CONCLUSIONS

FALCON-meta enables fast inference of metagenomic composition of ancestral whole genome raw data without overestimation. FALCON-meta measures relative similarity using linear time complexity. The identification of the composition is efficiently detected, enabling further classification of ancestral or contamination genomes and, further, genome reconstruction in the ancestral case. Overall, the Neanderthal, Denisova and Mammoth examples show that the method is robust and suitable for the analysis of aDNA from distinct sources. In one of the example we found and autheticated an ancient *Pseudomonas* bacteria in a Mammoth mitogenome.

## ACKNOWLEDGMENTS

We thank João Zilhão, Martin Kircher, Steven Salzberg, Svante Pääbo, David Reich and Gabriel Renaud for helpful comments and explanations, and Cláudio Teixeira for computational infrastructures. This work was partially funded by FEDER (Programa Operacional Factores de Competitividade - COMPETE) and by National Funds through the FCT - Foundation for Science and Technology, in the context of the projects UID/CEC/00127/2013, UID/BIM/04501/2013, UID/AMB/50017, PTCD/EEI-SII/6608/2014. The grant SFRH/BPD/111148/2015 to RMS and SFRH/BPD/77900/2011 to TC.

## AUTHOR CONTRIBUTIONS

D.P., A.P. and J.M.R. designed the algorithms. D.P. and M.H. implemented and tested the software. D.P., A.P., R.M.S., J.M.R and P.J.F. designed the experiments and D.P., A.P., R.M.S., J.M.R., M.H., T.C., P.J.F. analyzed the results. All the authors have written the manuscript.

